# Real-time Four-dimensional Imaging and Flow Dynamics Analysis of Cilia-driven Transport in Mammalian Tissues

**DOI:** 10.64898/2026.06.01.729054

**Authors:** Giancarlo Porcella, Ayse Tugce Sahin, Jens Keller, Janna C. Nawroth

## Abstract

Quantifying cilia-driven transport in mammalian tissues requires resolving rapid ciliary dynamics while preserving native three-dimensional geometry. Here we combine real-time four-dimensional light-field microscopy with tomographic particle tracking and physics-informed volumetric flow reconstruction to simultaneously image ciliary activity and reconstruct three-dimensional velocity fields in minimally dissected tissues. Applied across brain, respiratory, and reproductive epithelia, the workflow reveals conserved flow structures across individual and enables quantitative analysis of flow–tissue coupling.

## Introduction

Multiciliated epithelia in mammals generate internal fluid flows essential for diverse physiological processes, including cerebrospinal fluid (CSF) circulation^1^, mucociliary clearance^2^, and gamete transport^3^. Although motile cilia share conserved structural and biomechanical features, the resulting transport dynamics differ markedly across organs and are shaped by cilia organization, beat kinematics, organ geometry, and fluid properties. Recent studies have advanced our understanding of cilia-driven transport using flattened preparations or highly symmetric non-mammalian systems^2,4–8^. For example, flattened ex vivo brain ventricular preparations previously revealed spatially organized CSF flow microdomains generated by ependymal ciliary beating^4^. However, these surface-projected measurements cannot resolve how flows originating from opposing and non-coplanar ventricular walls interact within the native three-dimensional lumen. Measuring cilia-driven flow in preserved three-dimensional cavities of mammalian organs remains challenging due to the lack of high-speed imaging approaches capable of simultaneously capturing volumetric ciliary dynamics and fluid motion. Consequently, the organ-level organization of cilia-driven transport, its dependence on native organ architecture, and its conservation across individuals remain poorly understood. Recent computational studies suggest that Lagrangian transport analyses can reveal barriers, compartmentalization and residence dynamics in three-dimensional organs, but these approaches remain constrained by assumed ciliary forcing due to limited experimental data on ciliary beat patterns in mammalian tissues^9^. Here, we combine real-time volumetric imaging, tomographic particle tracking velocimetry and physics-informed flow reconstruction to measure cilia-driven transport directly in three dimensions. Using this newly conceptualized workflow, we identify recurrent flow structures across individuals, allowing us to generate a flow atlas of the mammalian brain ventricle. We further demonstrate quantitative flow–tissue coupling and present generalization of the method using airway and reproductive epithelia.

## Results

We first investigated the mouse third ventricle (3V), a ciliated brain cavity in which opposing and non-coplanar ventricular walls generate interacting cerebrospinal fluid (CSF) flows. Previous surface-projected measurements identified complex wall-adjacent flow networks^4^, but could not resolve how these flows interact within the native three-dimensional lumen. To simultaneously resolve ciliary dynamics, ventricular geometry, and fluid transport, we applied high-speed volumetric light-field microscopy using a commercially available imaging platform to minimally invasive wholemount ventricular preparations (**Fig.1a**). Samples were live-stained with tomato lectin to visualize cilia and seeded with 1 μm fluorescent microbeads to track fluid motion (Methods), enabling their simultaneous volumetric imaging at 40 volumes per second (vps). Direct visualization of three-dimensional particle trajectories, however, produced dense overlapping trajectory fields that limited quantitative interpretation (**Fig. 1b**). To enable quantitative flow analysis, tomographic particle tracking velocimetry (tPTV) was used to reconstruct three-dimensional particle trajectories and obtain divergence-free velocity fields within the imaged volume, through physics-informed radial basis function interpolation, using commercial software (**Fig. 1c, d**). This workflow reconstructs continuous volumetric velocity fields that directly relate flow organization to ventricular geometry and ciliary architecture, without inferring 3D flow from planar measurements^10,11^.

**Fig. 1.**
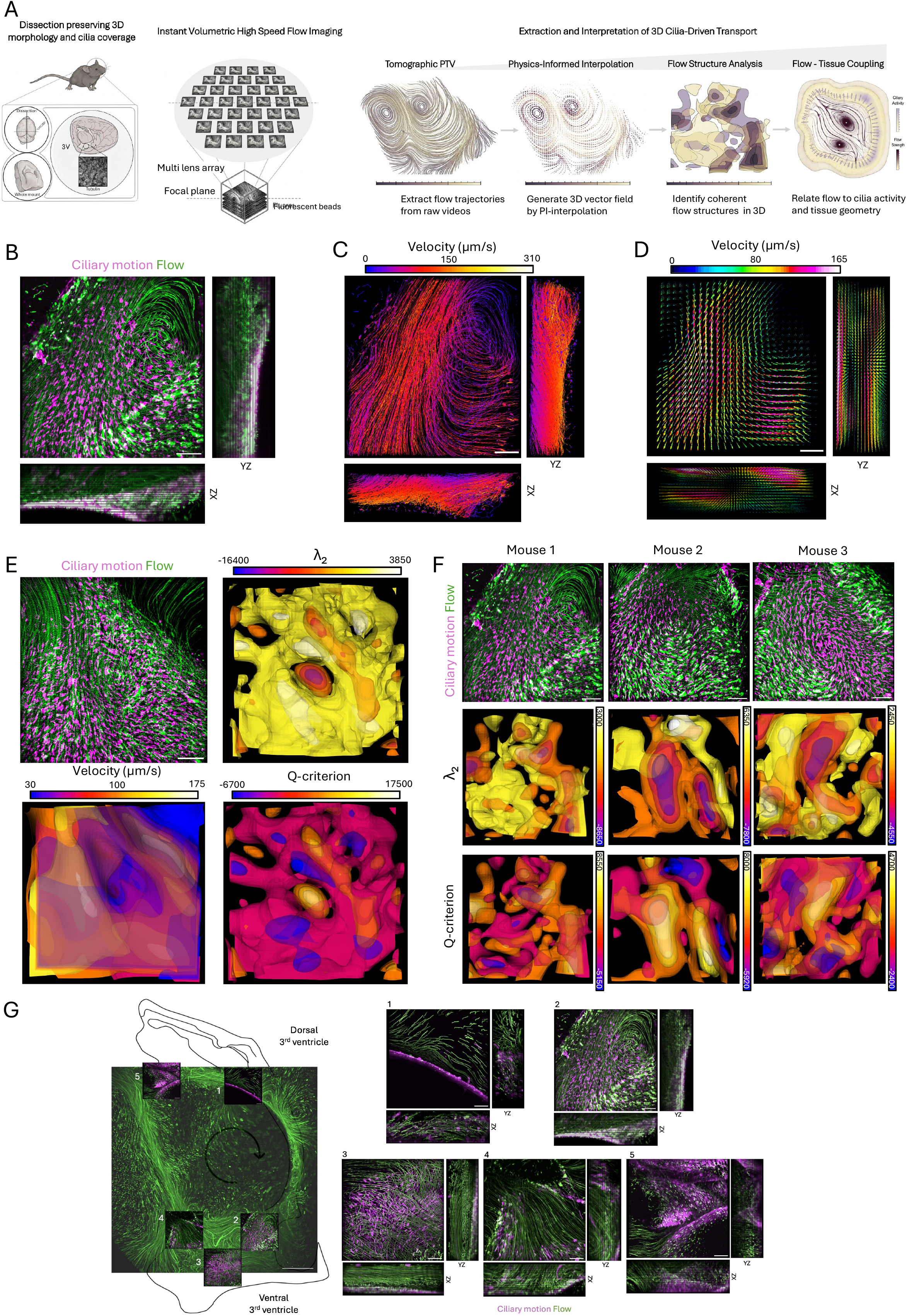
Real-time volumetric flow recording and characterization in the mouse brain ventricle. **A**, Schematic of the experimental workflow, from wholemount preparation to four-dimensional (4D) imaging and volumetric flow analysis. **B**, Planar and orthogonal views of the standard deviation projection of raw particle trajectories and ciliary beat motion at the entry of the ventral third ventricle (v3V), illustrating the complexity of three-dimensional flow organization. A direct visualization of this kind is not sufficient to fully interpret the complex 3D flow. **C**, Tomographic particle tracking velocimetry (tPTV) of the same region using a Lagrangian four-frame tracking approach; particle trajectories are color-coded by speed. **D**, Time-averaged velocity field obtained by physics-informed interpolation (PI-RBF) of tPTV data; vectors are color-coded by magnitude. **E**, Ciliary beat motion projection and three-dimensional vortex identification using Q-criterion and λ_2_ methods in the mouse v3V. Regions of high Q and low λ_2_ values define vortex cores. Isosurface rendering of velocity magnitude (vector length) further highlights a low-speed (high stagnation) region corresponding to the vortex core. **F**, ciliary beat motion projection (top) and Q-criterion and λ_2_ methods (bottom) applied to recordings of the same ventricular region in N=3 animals. **G**, Global flow atlas in the mouse 3V as highlighted by recurrent cilia configurations and corresponding conserved flow structures (scalebar = 500 µm); proceeding caudal-to-rostral, the flow descends from dorsal 3V (d3V) to the v3V (1), undergoes mixing through a vortex structure (2), reaches a splay flow convergence in the inner v3V (3), is driven upwards through the v3V outlet (4) and finally reaches the intersection between d3V and the fourth ventricle (5). Scalebars = 50 µm.

Application of Q-criterion and λ_2_ vortex analysis to the reconstructed volumetric velocity fields resolved three-dimensional vortex cores within the ventricular cavity (**Fig. 1e**, see Methods). Visualization of particle velocity magnitude, computed from frame-to-frame displacements and represented as vector length, further delineated a low-velocity region co-localizing with the vortex core. Comparative analysis across animals revealed recurrent and spatially conserved flow domains, including a recurrent vortex located at the flow entry in the ventral 3V (v3V) (**Fig. 1f**). Of note, while conventional volumetric imaging using sequential scans or z-stacks may reveal flow structures across multiple planes qualitatively, it cannot recover the time-resolved three-dimensional velocity vector field acquired by our workflow, including out-of-plane velocity components required for volumetric vortex identification and future transport-level analyses (**Extended Data Fig. 1**).

After establishing conserved local flow organization across animals, we assembled a global flow atlas of the ventral third ventricle, revealing recurrent flow patterns and pathways throughout the cavity (**Fig. 1g**, Supplementary Video 1).

To assess whether the workflow generalizes beyond the ventricular system, we extended it to additional mouse ciliated organs, namely the trachea and the oviduct, using minimally disruptive dissections that preserve relevant three-dimensional epithelial geometry, and the same cilia and flow labeling strategy used in the brain ventricles (**Fig. 2a, b**). In the trachea, where cilia-driven flow is predominantly oriented along the distal–proximal axis to support mucociliary clearance^12^, volumetric imaging resolves this directed transport within the tubular geometry (**Fig. 2c**). Similarly, imaging of the intact oviduct infundibulum following detachment from the ovary reveals organized flow directed toward the oviductal opening, driven by the coordinated activity of densely ciliated fimbriae (**Fig. 2d**). Cilia-driven flow and its functional implications remain only marginally characterized in female reproductive organs^3^, highlighting the utility of our workflow for understudied cilia-driven transport systems.

**Fig. 2.**
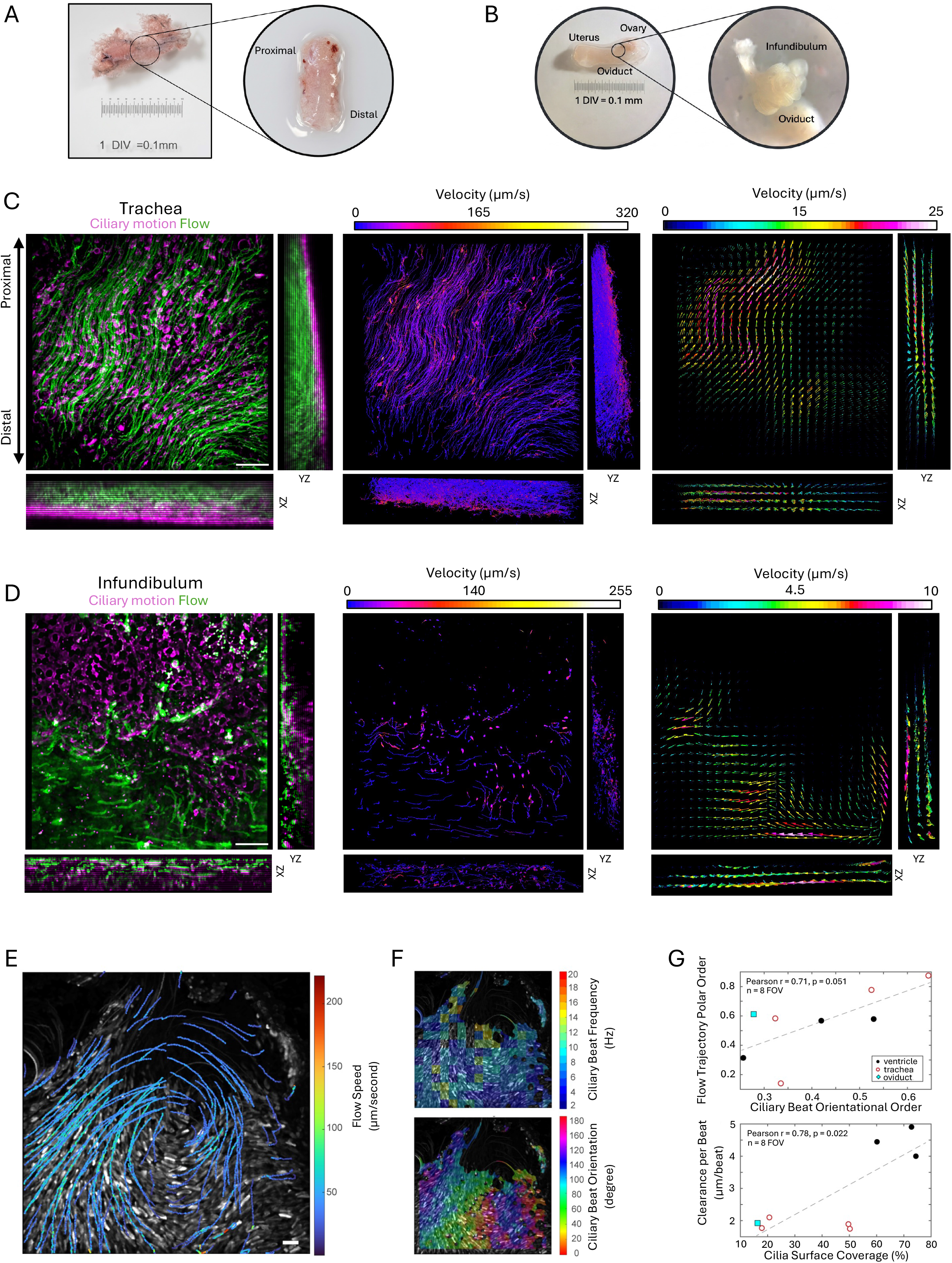
Volumetric recording and analysis of flow–tissue coupling across tissues. **A**, Overview of whole-mount isolation of the mouse trachea and **B**, the oviduct infundibulum. Dissections are performed to minimally disrupt native 3D morphology. **C**, Planar and orthogonal views of a mouse tracheal section (left to right): standard deviation projection of raw particle trajectories and ciliary beat motion; tPTV analysis of track speed; time-averaged velocity field obtained using PI-RBF interpolation. The analysis reveals overall upward flow characterized by multiple bends. **D**, Planar and orthogonal views of the mouse infundibulum opening (left to right): standard deviation projection of raw particle trajectories and ciliary beat motion; tPTV analysis of track speed; time-averaged velocity field obtained using PI-RBF interpolation. A slow, steady flow is observed in proximity to the ciliated fimbriae. Scalebars = 50 µm. **E**, Overlay of ciliary beat activity with surface-near flow trajectories color-coded by speed (scale bar = 20 µm). **F**, Corresponding spatial maps depicting ciliary beat frequency (top) and orientation (bottom). **G**, Quantitative analysis of the correlation between flow trajectory polar order and ciliary beat orientation (top), and between clearance per beat and ciliated coverage (bottom) in mouse 3V, trachea and oviduct (dots represent fields of view; data from n = 3 animals).

Because ciliary activity and fluid motion were recorded simultaneously across all tissues, the datasets further enabled direct investigation of flow–tissue coupling, i.e., the reconstructed flow fields could be directly related to local ciliary organization. In the same field of view, overlays of ciliary beat activity with speed-coded particle trajectories revealed spatial alignment between ciliary actuation and flow paths (**Fig. 2e**). Local maps of ciliary beat frequency and beat orientation (**Fig. 2f**) further enabled quantitative comparison of ciliary organization with flow output. Across fields of view and different ciliated organs, ciliary beat orientational order showed a positive trend with flow trajectory polar order, and cilia coverage was positively associated with clearance per beat, consistent with the expected flow–tissue coupling relationships (**Fig. 2g**), illustrating how the workflow can quantify flow–tissue coupling across ciliated tissues.

## Conclusion

Here, we combine minimally disruptive tissue preparation, high-speed volumetric light-field microscopy, and physics-informed flow reconstruction to enable simultaneous characterization of ciliary activity and fluid transport while preserving native three-dimensional tissue architecture. In contrast to planar or sequential plane-scanning approaches, the workflow reconstructs time-resolved volumetric velocity fields and resolves three-dimensional flow structures directly within preserved geometries. Application of the framework across mammalian ciliated organs revealed recurrent, reproducible, and spatially conserved flow organization between individuals, enabling quantitative comparison of flow structure, directionality, and flow–tissue coupling across tissues and experimental conditions. The workflow is furthermore compatible with complementary fluorescence-based modalities, including calcium imaging, enabling future interrogation of dynamic interactions between flow and cellular signaling.

Current limitations include the optical accessibility requirements of the sample preparation, finite imaging depth, and the inability to capture additional in vivo contributors to internal fluid dynamics, including secretion, pulsatility, and body movement. Nevertheless, the framework establishes a generalizable approach for analyzing cilia-driven transport in preserved three-dimensional tissues and provides direct access to ciliary beat dynamics and three-dimensional flow organization that is inaccessible to conventional two-dimensional imaging strategies.

## Acknowledgments

This work was funded by the European Research Council (ERC: ERC-STG 950219 and ERC-POC 101212905, J.N.) and Volkswagen Foundation (Pioneering Research Award 9D174, J.N.). We thank ZEISS (Carl Zeiss Microscopy GmbH) for access to Lightfield4D technology and technical support. We also thank Prof. Dr. Ali Önder Yildirim (Lung Health Institute, Helmholtz Munich) and the German Mouse Clinic (Helmholtz Munich) for providing mouse samples.

## Online Methods

### Real-time volumetric light-field microscopy

The ZEISS LSM 990 equipped with the Lightfield 4D module enables real-time volumetric imaging by placing a microlens array between the objective and detector, thereby capturing multiple angular views of the emitted signal in a single acquisition. These multiperspective images were reconstructed by deconvolution into time-resolved volumetric image stacks, allowing acquisition at up to 80 reconstructed volumes per second (vps).

### Volumetric light-field imaging of ciliary beat and particle flow

To record both ciliary motion and the resulting flow patterns on the highly irregular, complex 3D morphology of the tissue sections, samples were placed face-down in a 35 mm glass-bottom dish (ibidi cat. no. 81218) and immersed in 3 ml Leibovitz L-15 medium (Thermo Fisher Scientific, 11415064) at room temperature, taking care to maintain local tissue geometry and ciliary activity. Cilia were live-stained with Lycopersicon esculentum (Tomato) Lectin DyLight 649 (Thermo Fisher Scientific, L32472), diluted 1:500 in L-15 medium (brain ventricles and infundibulum) or fluorescent-dye conjugated wheat germ agglutinin (Thermo Fisher Scientific; W11261) diluted 1:100 in L-15 medium (tracheas). Cilia-generated flow was visualized by adding 1 µm carboxylate-modified green- (brain ventricles and infundibulum) or red- (tracheas) fluorescent microbeads (Thermo Fisher Scientific, F8819/F8803) diluted 1:200 in L-15 medium.

For ciliary beat frequency (CBF) quantification, ciliary beat was acquired at 80 vps for 3 s with a water-immersion 40× objective to record the beating trajectories of cilia bundles. To simultaneously acquire ciliary motion and the associated flow patterns, movies were recorded at 40 vps with 40× and 10× objectives for 7.5 s, allowing the direct superposition of the two signals.

### Animal studies

All animal tissue samples were collected immediately after euthanasia by collaborating laboratories at the Helmholtz Zentrum München, whose experimental procedures were approved by the local authority and carried out in accordance with the European Directive 2010/63/EU on the protection of animals used for scientific purposes, the German Animal Welfare Act (Tierschutzgesetz, TierSchG), and the German Animal Welfare Experimental Animal Ordinance (Tierschutz-Versuchstierverordnung, TierSchVersV). For all experimental procedures, adult wild type C57BL/6N mice of both sexes between P90-P300 were used.

### Tissue preparation for standard and 4D light-field imaging

All tissues were processed immediately after euthanasia and maintained in ice-cold Hanks’ Balanced Salt Solution (HBSS, without phenol red) throughout dissection to preserve ciliary activity. Samples were imaged within hours of preparation unless otherwise stated. Under these conditions, ciliary motility in brain ventricular preparations remained stable for up to 48 h when stored at 4 °C.

#### Brain ventricles

Mouse brains were rapidly extracted from the skull and transferred ventral side up into a PDMS-coated dissection dish filled with ice-cold HBSS. During the extraction, particular care was taken to avoid mechanical perturbation of the ventral brain surface, which lies in close proximity to the third ventricle (3V). The olfactory bulbs and cerebellum were removed to facilitate access. The hemispheres were stabilized using fine cactus pins placed laterally and gently separated along the midline using forceps while carefully severing connecting structures, including the anterior commissure and thalamic regions. Each hemisphere was subsequently trimmed to obtain ∼1–1.5 mm-thick sections exposing the 3V wall. The resulting semi-intact wholemounts, preserving the native curvature and organization of the ventricular surface, were transferred to 35 mm glass-bottom dishes and oriented with the ciliated surface facing the objective for imaging.

#### Infundibulum

The reproductive tract was isolated from female mice by excising both uterine horns at the uterine junction while preserving the ovaries and surrounding adipose tissue. Samples were maintained in ice-cold HBSS until further processing. Dissections were performed by anchoring the distal uterine horn and ovarian fat pad and gently separating the ovary to expose the infundibulum without disrupting its three-dimensional structure. Excess adipose and connective tissue were carefully removed. The oviduct was severed at the uterotubal junction to isolate the coiled oviduct with an intact infundibulum. Samples were transferred to glass-bottom dishes and oriented to expose the fimbrial surface toward the imaging plane, preserving local 3D geometry.

#### Tracheas

The trachea was exposed via a ventral midline incision extending from thorax to jaw. Segments of approximately 5 mm were excised and transferred to ice-cold HBSS. Surrounding connective tissue and fat were removed under a stereomicroscope. To expose the ciliated luminal surface, tracheal segments were opened longitudinally along the dorsal side and placed epithelial side down in glass-bottom dishes. Samples were submerged in HBSS and gently flattened using a coverslip secured with small amounts of silicone grease at the corners to decrease curvature while preserving epithelial integrity.

### Data processing and analysis

#### Qualitative visualization of ciliary beat and flow

Ciliary motion maps were generated using the temporal standard deviation projection of the cilia channel. Qualitative flow patterns were visualized using the temporal maximal projection of the microbead channel.

#### Light-field volume reconstruction

Raw light-field image volumes were first reconstructed and converted into volumetric image stacks compatible with the DynamicStudio workflow using a custom Python script, which reformats LightField4D output into a three-dimensional time-resolved dataset. The resulting volumes were subsequently pre-processed to enhance signal-to-noise ratio and particle detectability.

#### Particle detection and tomographic PTV

Three-dimensional flow reconstruction was performed using Dantec Dynamics DynamicStudio (version 8.6), following established workflows for tomographic particle tracking velocimetry (Tomographic PTV)^13^. Fluorescent tracer particles (1 μm) were identified in three dimensions using intensity-based segmentation and tracked over time using a 4-frame tracking algorithm, resulting in discrete Lagrangian particle trajectories.

#### Physics-informed interpolation to volumetric velocity fields

The Lagrangian particle trajectories were used to reconstruct continuous volumetric velocity fields. To this end, a physics-informed interpolation approach based on radial basis functions was applied^14–16^, enforcing incompressibility constraints to obtain divergence-free velocity fields while preserving experimentally measured particle motion. This procedure yields time-resolved three-dimensional velocity vector fields across the imaged volume, enabling quantitative characterization of flow organization in relation to tissue geometry and ciliary activity. Reconstruction parameters, including particle density thresholds and tracking tolerances, were optimized to ensure robustness and reproducibility across samples while minimizing artifacts arising from tracking uncertainty or sparse sampling.

#### Flow-structure analysis

Flow-structure analysis was performed on the reconstructed time-averaged volumetric velocity fields. Spatial velocity gradients were computed from the interpolated velocity fields and used to calculate Q-criterion and λ_2_ vortex metrics. Here, *S* and Ω denote the symmetric and antisymmetric components of the velocity-gradient tensor, respectively. The Q-criterion identifies regions in which local rotation dominates strain, with 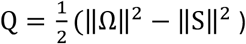, whereas λ_2_ analysis identifies vortex cores where the second-largest eigenvalue of *S*^2^ + Ω^2^ is negative. Isosurfaces and maps were generated using fixed thresholds applied consistently across comparable fields of view.

#### Flow metrics near ciliated surface

Flow metrics near the ciliated surface were quantified from maximum-intensity projections of the flow channel to enable direct comparison with projected ciliary activity maps. Flow trajectories were detected using ImageJ/Fiji plugin TrackMate^17^. The average flow speed for each field of view was defined as the mean of the median speed of all accepted trajectories. Flow trajectory polar order was calculated from the distribution of trajectory directions as *P*_*f*_ = | ⟨*e*^*iϕ*^ ⟩ |, where *ϕ* denotes the mean trajectory direction.

#### Ciliary beat analysis

Ciliary beat analysis was performed from maximum-intensity projections of the cilia channel. Ciliary beat frequency, coverage, and orientation were automatically computed using MATLAB-based Fourier analysis, thresholding of detected motion, and steerable wavelet analysis, respectively. Active cilia coverage was defined as the fraction of pixels with detected ciliary motion. Ciliary beat orientational order was calculated as 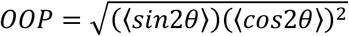, where *θ* denotes local ciliary beat orientation; (OOP = 0) indicates randomly oriented beating and (OOP = 1) indicates aligned beating^2^.

#### Flow–tissue coupling analysis

Flow–tissue coupling was assessed by comparing ciliary organization metrics with flow output metrics across fields of view^2^. Clearance per beat was defined as average flow speed divided by average ciliary beat frequency, yielding the average tracer displacement per beat cycle. Associations between ciliary beat orientational order and flow trajectory polar order, and between active cilia coverage and clearance per beat, were assessed across ventricular, tracheal and oviduct datasets. Linear associations between ciliary organization and flow output metrics were assessed using Pearson correlation, based on the expected locally linear relationships between these quantities^2^.

### Reproducibility and quality-control

A minimum of three animals were analyzed for each tissue type. For each animal, a minimum of three fields of view were acquired to ensure reproducibility of both ciliary dynamics and the resulting flow patterns, and to account for biological and spatial heterogeneity within samples. For brain ventricular preparations, imaging regions were selected to represent distinct anatomical domains of the ventricular system associated with characteristic flow patterns. The highly conserved morphology of the mouse third ventricle enabled reliable identification and alignment of corresponding regions across animals, thereby facilitating cross-sample comparisons.

To ensure data quality, only samples exhibiting preserved tissue architecture and sustained ciliary activity were included in the analysis. Preparations were imaged within 30 min after dissection, and ciliary motility was routinely verified prior to acquisition. Imaging conditions, including labeling, bead concentration, and acquisition parameters, were kept constant across experiments.

For volumetric flow reconstruction, particle detection thresholds and tracking parameters were optimized and then held constant across comparable datasets: search volume (x,y,z): 6 µm x 6 µm x 3 µm; minimum particle volume (voxels):7; minimum steps tracked per particle: 4. Trajectories with insufficient temporal continuity or ambiguous particle assignment were excluded based on these thresholds. The density of successfully tracked particles was compared to raw 3D particle recordings to ensure adequate spatial sampling for reliable interpolation. The physics-informed reconstruction enforcing incompressibility provided an additional constraint to reduce noise and mitigate tracking artifacts. Derived flow quantities (including velocity fields and vortex metrics) were computed using standardized parameter settings and threshold values applied uniformly across datasets, enabling reproducible identification and comparison of flow structures.

## References

1. Fame, R. M., Cortés-Campos, C. & Sive, H. L. Brain Ventricular System and Cerebrospinal Fluid Development and Function: Light at the End of the Tube: A Primer with Latest Insights. BioEssays News Rev. Mol. Cell. Dev. Biol. 42, e1900186 (2020).

2. Roth, D. et al.. Structure and function relationships of mucociliary clearance in human and rat airways. Nat. Commun. 16, 2446 (2025).

3. Ezzati, M., Djahanbakhch, O., Arian, S. & Carr, B. R. Tubal transport of gametes and embryos: a review of physiology and pathophysiology. J. Assist. Reprod. Genet. 31, 1337–1347 (2014).

4. Faubel, R., Westendorf, C., Bodenschatz, E. & Eichele, G. Cilia-based flow network in the brain ventricles. Science 353, 176–178 (2016).

5. Ling, F. et al.. Flow Physics Explains Morphological Diversity of Ciliated Organs. BioRxiv Prepr. Serv. Biol. 2023.02.12.528181 (2024) doi:10.1101/2023.02.12.528181.

6. Yuan, S. et al.. Motile cilia of the male reproductive system require miR-34/miR-449 for development and function to generate luminal turbulence. Proc. Natl. Acad. Sci. 116, 3584–3593 (2019).

7. Ramirez-San Juan, G. R. et al. Multi-scale spatial heterogeneity enhances particle clearance in airway ciliary arrays. Nat. Phys. 16, 958–964 (2020).

8. Ling, F. et al.. Flow physics guides morphology of ciliated organs. Nat. Phys. 1–8 (2024) doi:10.1038/s41567-024-02591-0.

9. Herlyng, H. & Shadden, S. C. Characterization of coherent flow structures in brain ventricles. Preprint at 10.48550/ARXIV.2603.18849 (2026).

10. Siyahhan, B. et al.. Flow induced by ependymal cilia dominates near-wall cerebrospinal fluid dynamics in the lateral ventricles. J. R. Soc. Interface 11, 20131189 (2014).

11. Yoshida, H. et al.. Effect of cilia-induced surface velocity on cerebrospinal fluid exchange in the lateral ventricles. J. R. Soc. Interface 19, 20220321 (2022).

12. Bustamante-Marin, X. M. & Ostrowski, L. E. Cilia and Mucociliary Clearance. Cold Spring Harb. Perspect. Biol. 9, a028241 (2017).

13. Kitzhofer, J., Brücker, C. & Pust, O. Tomo PTV using 3D Scanning Illumination and Telecentric Imaging. 8th International Symposium On Particle Image Velocimetry. Melbourne, Victoria, Australia, August 25–28, 2009

14. Sperotto, P., Watz, B. & Hess, D. Meshless track assimilation (MTA) of 3D PTV data. Meas. Sci. Technol. 35, 086005 (2024).

15. Sperotto, P., Hessa, D. & Ergina, F. G. 3D Tomographic PTV investigation of three leap frogging vortex rings. 1st European Fluid Dynamics Conference (EFDC1). 16 ™20 September 2024, Aachen, Germany

16. Sperotto, P., Petersson, P., Watz, B. & Hess, D. Instantaneous 3D Flow Field Reconstruction with Physics Informed Radial Basis Functions. 15th International Symposium on Particle Image Velocimetry – ISPIV 2023. June 19–21, 2023, San Diego, California, USA

17. Tinevez, J.-Y. et al.. TrackMate: An open and extensible platform for single-particle tracking. Methods 115, 80–90 (2017).

